# The Dynamics of Resting-State Alpha Oscillations Predict Individual Differences in Creativity

**DOI:** 10.1101/818229

**Authors:** Naomi Prent, Dirk J.A. Smit

## Abstract

The neuronal mechanisms underlying creativity are poorly understood. Here, we investigated whether temporal dynamics of functional brain activity is a biomarker of creative ideation. Specifically, we investigated whether long-range temporal correlations in fluctuating resting-state alpha oscillations predict human creativity. Because lower temporal correlations reflect faster brain state switching, and faster state switching may be associated with increased flexibility of mind, we hypothesized that subjects with lower temporal correlations would show higher creativity. Creativity was measured by self-rating, examiner-rating and the alternative uses task in 40 healthy young adults, and scored on dimensions of verbal fluency, originality, elaboration, usefulness, and flexibility. For each dimension, the total number of subject-reported alternative uses that matched the criterium was noted (the quantity measure), as well as the proportion of uses that matched the dimensional criterium. A principal components analysis confirmed the two-component structure of quantity and quality. Eyes-open resting-state brain activity was measured by electroencephalogram (EEG) with 128 channels. Scaling exponents β were derived from spectral analysis of the amplitude modulation of 8-13 Hz oscillations, where high exponents β reflect lower decay of autocorrelation and slower switching of brain. Partial correlation analysis was used controlling for gender and age, and a cluster permutation test was performed correcting for multiple testing. Significant negative correlations between creativity and temporal correlations were observed, most prominent in right central/temporal brain areas. To our knowledge, this is the first demonstration that individual variation in the intrinsic dynamics of the brain may offer a neuronal explanation for individual variation in both the quality and quantity of creative ideation.

---

Creativity plays an important role in almost every field of life: in art, music, science, economy, and technology. Creativity is our ability to change existing thinking patterns, flexibly break with ongoing trains of thought, and make innovation possible (Dietrich & Kanso, 2010). Sternberg and Lubart (1996) defined creativity as ‘the ability to produce work that is both novel and appropriate’. And according to Guilford (1950), creativity is divergent thinking in which originality, flexibility, and fluency are the main components. Creativity correlates with well-being and adaptive behavior, which has led to a large body of investigations into its neurobiological underpinnings. Recent reviews are increasingly converging on brain areas and networks involved in creative ideation and explaining individual differences in creativity (Dietrich and Kanso, 2010; Beaty et al., 2016; Khalil et al., 2019). Relatively under-investigated is how these brain areas affect creativity and the flexibility of thought it requires. Here, we investigate whether temporal dynamics in functional brain activity indexes the propensity to switch brain states, thus increasing flexibility in thought as one of the key components in creative ideation.

Creative abilities may have been selected for by an evolutionary advantage offering advantages for survival by inventing solutions for life threatening situations, but also by enhancing social cohesion by music and dance. It has been observed that creative people have more sexual partners than average (Thys, Sabbe & De Hert, 2012). On the downside, creativity may be enhanced in specific mental disorders such as schizophrenia, and the liability for the disease appears correlated with creative behaviors (Post, 1994; Kyaga et al., 2011; Power et al., 2015) suggesting that creativity and these mental disorders share a genetic background. However, those genetically at risk for these disorders, but phenotypically unaffected, may show increased procreation rates, which may explain why genetic risk variants are maintained in the population despite the reduced rates of procreation in affected individuals (Van Dongen & Boomsma, 2013).

Reviews of investigations into the neuronal mechanisms underlying creativity conclude that they are still poorly understood (Dietrich and Kanso, 2010). Creative thinking in a does not appear to depend on any single mental process or brain region (Dietrich and Kanso, 2010; Stevens and Zabelina, 2019). Idea generation begins with retrieval of common and old ideas, followed by processes of mental simulation and imagination to the actual generation of novel and creative ideas (Schwab et al., 2014). Consequently, it may be assumed that multiple specific neurocognitive processes at multiple brain areas are associated with different types and stages of creativity. According to Dietrich and Kanso (2010), this could be an explanation for highly variegated results of research on brain regions related to creativity.

The most consistent finding in the neuroscientific study of creativity seems to be the increased power of 10 Hz (alpha) oscillations during creative ideation (Martindale and Mines, 1975; Martindale and Hasenfus, 1978; Fink and Benedek, 2014; Luft et al., 2018; Stevens and Zabelina, 2019), although some have argued that the results are inconclusive (Dietrich and Kanso, 2010). Further EEG studies suggested that generation of original ideas is associated with alpha synchronization in the prefrontal cortex (Fink et al., 2011) and in the parietal and temporal sites of the right hemisphere (Schwab et al., 2014), and that creative persons show stronger frontal activity, whereas lower creative individuals show increased parietal activity, suggests that lower and higher creative individuals may use different strategies of divergent thinking (Jauk et al., 2012). In addition, neuroimaging studies revealed strong activation in prefrontal regions and parietal cortex of the left hemisphere (Fink et al., 2009; Dietrich and Kanso, 2010; Benedek et al., 2014). A meta-analysis suggested right-hemisphere dominance for specific creative tasks (Mihov et al., 2010).

The nature of creative thinking, specifically, being able to produce thoughts that are novel (Runco and Jaeger, 2012), may partially depend on the ability of the brain to produce activity patterns that are independent of previous states. Creative thinking depends on break from habitual thinking and associations (Luft et al., 2018), which might depend on a decorrelation in the temporal dynamics of brain activity patterns. Research has shown that spontaneous brain activity fluctuations (EEG neuronal oscillations, microstates, fMRI) show substantial temporal correlations (Linkenkaer-Hansen et al., 2001; Miller et al., 2009; Ville et al., 2010; Palva et al., 2013; Smit et al., 2013) with power-law decay of the autocorrelation. This results in power-spectra *P*(*f*) ∝ 1/*f*^*−β*^, where *P* is power, *f* is frequency and *β* is the power-law exponent (He, 2014). In double-log PSD plots *β* represents the steepness of the downward slope generally observed. Positive values of *β* indicate larger power for slow frequencies relative to higher frequencies, resulting in the signal remaining in a similar state for prolonged times. For example, the modulation of alpha band oscillations showed significantly nonzero values. Moreover, it was observed that individuals show stable and heritable variation in the autocorrelation decay parameter (Linkenkaer-Hansen et al., 2007; Smit et al., 2010).

Previous studies have shown that Interindividual variability in the temporal correlations in brain dynamics have been found to predict performance in simple cognitive behavior like finger tapping and the detection of threshold stimuli (Smit et al., 2013; Palva et al., 2013). It is unknown whether brain dynamics may also have an effect on more complex behavior like creativity. Therefore, our aim was to investigate whether temporal correlations in resting-state alpha oscillations are a biomarker of human creativity. Since alpha oscillations have been found to be involved in the suppression of habitual, and lead to more creative solutions (Luft et al., 2018), we hypothesized that a faster decay in autocorrelation of alpha power would result in increased creative output.

## Methods

### Participants

Participants were recruited within the VU University community. Participants were selected on their age (18-30 years), individuals using antipsychotics, sedative drugs, or psychoactive medication like Ritalin, having experienced insomnia the past 3 days, epilepsy, or having ever experienced unconsciousness for more than 5 minutes were excluded. All participants were informed about the nature of the research and signed an informed consent. The study was approved by the ethical review committee of the Faculty of Psychology and Education of the VU University Amsterdam.

The sample consisted of 40 participants (20 males), with ages ranging from 20 to 27 years, and a mean age of 22.55 (*SD*=1.77). The majority of the participants (87.5%) were born in the Netherlands, others in Belgium, Russia, France or Iran. Furthermore, 22.5% of the participants had one or both parents born outside Europe. At the time of the research, more than half of the participants followed a university education (57.5%) and others followed college education (20%), high school (7.5%) or did no education (15%). The participants had different educational backgrounds: Behavioral and Social Sciences (30%), Natural Sciences (20%), Health and Movement (15%), Economics, Business and Law (12.5%), Computer Science, Mathematics and Business (12.5%), Art, Culture and History (5%) and Language and Communication (5%). None of the participants had ever experienced psychosis and two participants had a family member who had ever had a psychosis. 92.5% of the subjects were right-handed.

### Materials & Procedure

First, a digital survey with general questions (gender, age etc.) was administrated via Qualtrics (Qualtrics, Provo, UT, version 2015) on a computer. Participants were asked to rate their own creative ability on a 10-point scale. Next, Qualtrics was used for taking the Alternative Uses Task (AUT)(Guilford, 1967) for the measurement of divergent thinking. (Vosburg, 1998) reported an overall Cronbach’s alpha of .86 for the AUT. During the task, participants were asked to name as many unusual uses as possible for 4 various conventional, everyday objects (paperclip, newspaper, brick, and shoe) in 3 minutes per object.

Next, the participants were connected to the EEG system and the resting-state brain activity was measured. The subject was instructed to sit silently with eyes closed and minimize movement for three minutes, followed by a three-minute measurement with eyes open. This resting-state brain activity measurement was performed again after several minutes—after performing a Wisconsin Card Sorting Task—resulting in a total of six minutes eyes-open and 6 minutes eyes-closed. Only the eyes-open condition was used for the current analyses, consistent with previous investigations (Smit et al., 2013). Statistical Package for Social Scientists (IBM-SPSS Statistics, version 20, 2011) and MATLAB (The Mathworks Inc., Natick, MA, version 2014) were used for data analysis.

### EEG registration

The electrical brain activity was recorded by the 128 channel system BioSemi Ag/AgCl active electrode configuration. The active ground electrodes Common Mode Sensor (CMS) and passive ground electrode Driven Right Leg (DRL) formed a feedback loop, resulting in very high quality recordings even at high impedance contact. The sampling frequency was 1024 Hz and the low-pass filter during acquisition 208 Hz. The 128 electrodes were attached to a stretch lycra electrode cap placed on the head of the participant. SignaGel Electrode Gel conducted the signal of the scalp to the electrode. Furthermore, one electrode was placed 2 centimeters beneath each eye vertically aligned with pupil and one electrode 2 centimeters outside the outer canthi of each eye of the participant. These 4 electrodes recorded eye movements and blinks. Then, the electrodes were connected to the amplifier and computer and the functioning of the electrodes was checked.

### Scoring Alternative Uses Task

The responses of the Alternative Uses Task were scored on five main indices: Fluency, Flexibility, Originality, Elaboration and Usefulness. Fluency is the total number of ideas produced for each object. The valid number of ideas (Fluency-valid), with only unusual ideas, is also included. Flexibility is the number of generated categories of ideas for each object. Originality is the novelty of the ideas (statistically infrequent). Elaboration is the amount of detail of the responses and Usefulness is the functionality of the idea. The scoring of Fluency and Flexibility require simple counting, whereby every idea and every category are worth 1 point. The scoring of Originality is that ideas given by only 5% of the group count for 1 point (i.e., no more than 2 others gave the same answer were deemed original). In addition, responses rated as detailed received 1 point. For example, the answer ‘you can use a shoe to put in a cup so that the cup does not fall over in the car while driving’ was rated as detailed, because it provided more information than the use as a drink holder. Responses that were rated as functional will also get 1 point. The scores per object are added up to a total score for each category separately. In addition, because all these categories are based on the amount of responses (quantity), 3 ratios (number of points per category divided by total number of responses) for Originality, Elaboration and Usefulness were made and used as separate scores, for providing more information about the quality of the responses. Furthermore, quality was assessed by the examiner (N.P.), who divided the participants into 5 groups from least creative to most creative.

### EEG analysis

First, EEG signals from the 128 locations were preprocessed with a zero-phase bandpass FIR filter in the frequency bands of alpha (8–13 Hz) oscillations. Next, the envelope around the oscillations was determined as the absolute value of the complex valued |*s*(*t*) + *iH*(*s*(*t*))| of the filtered signal *s(t)* and its Hilbert transform 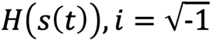. Figure 1 shows an example of a filtered signal with the amplitude envelope. We assessed a subject’s temporal stability in brain oscillatory alpha activity by calculating long-range temporal correlations of these modulating (waxing and waning) alpha oscillation strength. The temporal correlations were estimated using spectral analysis of the amplitude envelope signal. The power spectrum density estimated using an FFT with 64s box windows (50% overlap). Exponent *β* was defined as the absolute value of the slope of the linear regression of log-power on log-frequency from 0.0156 to 2 Hz. High levels of *β* indicate (relatively) high power for slow frequencies in the amplitude modulation, and therefore stronger temporal correlations, and more stability in the brain signals.

**Figure 1.**
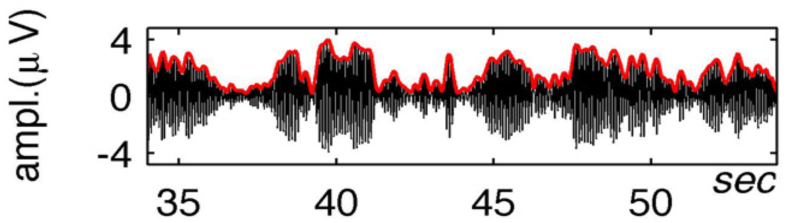
Illustration of brain oscillations filtered at 9 –13 Hz (black) and the amplitude envelope (red) determined using the Hilbert transformation (recreated from Smit et al., 2013). The highly clustered alpha oscillation bursts indicate high autocorrelation, strong temporal correlations and thus a high exponent beta.

A goodness-of-fit of linearity was estimated with *r*^2^ performed in the regression in the spectral analysis of the amplitude envelope.

### Statistics

First, the different creativity scores, which included Self-rating (SR), Examiner-rating (EX), Fluency (F), Fluency-valid (FV), Flexibility (FX), Originality and Originality-ratio (O and OR), Elaboration and Elaboration-ratio (E and ER), Usefulness and Usefulness-ratio (U and UR) were correlated to examine the convergent validity. Principle Component Analysis (PCA) was then used to reveal possible dimensions of creativity and to provide component scores. Finally, components were selected based on the eigenvalue distribution, rotated using promax rotation, and component scores calculated. These were correlated with the power-law exponents of the amplitude modulation of 8.0-13-0 Hz for each of the 128 electrodes measured during the eyes open condition. Standard partial correlation analysis was used controlling for gender and age.

A Monte Carlo permutation test was performed correcting for multiple testing, non-normal distributions, as well as outliers. In this approach, we kept the 128 scaling exponents (predictors) per subject intact, reshuffling the dependent data (quantitative creativity, qualitative creativity or total creativity). Next, we calculated the 128 correlations of these predictors with the residuals after regressing out the effects of age and gender from the creativity scores, and sorted the outcome. The three lowest (strongest negative) correlations were noted, and the next permutation performed. This resulted in null-distributions for the minimum and the two next minimal correlations, to which the actual correlations were compared using the 97.5 percentile.

## Results

### Validation of the creativity variables

To examine whether the different creativity variables (Self-rating, Examiner-rating and Alternative Uses Task scores) used in were related, standard Pearson correlation analysis was performed (Table 1). For completeness, means with standard deviations for men, women and the total sample are presented in Table 2. The correlations between the count-based scoring scales of the AUT are indicated by the dashed square (. Most of these are very strong, with the exception of Elaboration. The Self-rating variable of creativity correlates significantly with large effects with most indices of the AUT, moderate correlations were found in the correlation with Elaboration and Examiner-rating. This suggests that the Self-rating is primarily based on the number of ideas generated. Furthermore, the Examiner-rating shows medium-to-large correlations on all the categories of the AUT, but it additionally strongly correlates significantly with Originality ratio, Elaboration ratio and Usefulness ratio. This indicates that the ratios provide a good indication that a second source of variation may be found next to the count-based measures of Fluency, Fluency-valid, Flexibility, Originality, Elaboration, and Usefulness.

**Table 1:** Pearson correlation coefficients for the creativity scores (N=40).

|  | SR | EX | F | FV | FX | O | E | U | OR | ER | UR |
| --- | --- | --- | --- | --- | --- | --- | --- | --- | --- | --- | --- |
| Self-rating (SR) | 1 | **.48** | **.45** | **.51** | **.51** | **.41** | **.35** | **.52** | **.33** | .18 | .18 |
| Examiner-rating (EX) | **.48** | 1 | **.39** | **.61** | **.65** | **.60** | **.68** | **.74** | **.57** | **.42** | **.72** |
| Fluency (F) | **.45** | **.39** | 1 | **.94** | **.91** | **.70** | .20 | **.83** | .25 | -.26 | -.06 |
| Fluency-valid (FV) | **.51** | **.61** | **.94** | 1 | **.99** | **.78** | **.35** | **.95** | **.39** | -.13 | .24 |
| Flexibility (FX) | **.51** | **.65** | **.91** | **.99** | 1 | **.77** | **.39** | **.95** | **.42** | -.09 | .28 |
| Originality (O) | **.41** | **.60** | **.70** | **.78** | **.77** | 1 | **.43** | **.78** | **.81** | .04 | .27 |
| Elaboration (E) | **.35** | **.68** | .20 | **.35** | **.39** | **.43** | 1 | **.45** | **.42** | **.81** | **.40** |
| Usefulness (U) | **.52** | **.74** | **.83** | **.95** | **.95** | **.78** | **.45** | 1 | **.45** | -.01 | **.47** |
| Originality-ratio (OR) | **.33** | **.57** | .25 | **.39** | **.42** | **.81** | **.42** | **.45** | 1 | .26 | **.44** |
| Elaboration-ratio(ER) | .18 | **.42** | -.26 | -.13 | -.09 | .04 | **.81** | -.01 | .26 | 1 | **.37** |
| Usefulness-ratio (UR) | .18 | **.72** | -.06 | .24 | .28 | .27 | **.40** | **.47** | **.44** | **.37** | 1 |

*Note.* Nominally significant correlations ( $p < 0.05$ ) are indicated in bold.

**Table 2:** Means (M) with standard deviation (SD) for the creativity scores of the total sample, men (N=20) and women (N=20).

|  | Total |  | Men |  | Women |  |
| --- | --- | --- | --- | --- | --- | --- |
|  | *M* | *SD* | *M* | *SD* | *M* | *SD* |
| Self-rating | 6.73 | 1.69 | 6.60 | 1.90 | 6.85 | 1.50 |
| Examiner-rating | 3.05 | 1.06 | 3.01 | 1.00 | 3.05 | 1.15 |
| Fluency | 29.15 | 11.69 | 32.70 | 12.34 | 25.60 | 10.09 |
| Fluency-valid | 25.08 | 10.49 | 27.50 | 10.51 | 22.65 | 10.15 |
| Flexibility | 24.03 | 9.84 | 26.20 | 9.67 | 21.85 | 9.75 |
| Originality | 5.97 | 4.17 | 6.60 | 4.41 | 5.35 | 3.94 |
| Elaboration | 5.57 | 4.35 | 4.10 | 3.74 | 7.05 | 4.50 |
| Usefulness | 22.05 | 9.85 | 23.60 | 10.13 | 20.50 | 9.57 |
| Originality-ratio | .19 | .11 | .19 | .10 | .20 | .14 |
| Elaboration-ratio | .21 | .17 | .13 | .10 | .29 | .18 |
| Usefulness-ratio | .76 | .20 | .73 | .23 | .79 | .16 |

### PCA revealed a quantitative and qualitative component

The scoring procedure for the Alternative Uses task was based on previous research (Cropley, 2000; Runco & Acar, 2012; Vosburg, 1998). Many studies only counted for Fluency. However, Fluency is only one aspect of divergent thinking and does not guarantee the quality of ideas (Vosburg, 1998), and Fluency may not be as closely tied to creativity as Originality and Flexibility (Runco & Acar, 2012). Because some tests also included Elaboration (Runco & Acar, 2012), whereas others included Usefulness (Cropley, 2000; Vosburg, 1998), it was decided to include all these indices in addition to the rating scores.

Principal Component Analysis (PCA) revealed the presence of only 2 components with eigenvalues exceeding 1 (eigenvalue Component 1= 6.03, eigenvalue Component 2= 2.33), which corresponds with inflexion point at eigenvalue 3 (Figure 2). The two-component solution explained a total of 76.0% of the variance, with Component 1 contributing 54.8% and Component 2 contributing 21.2% respectively.

**Figure 2.**
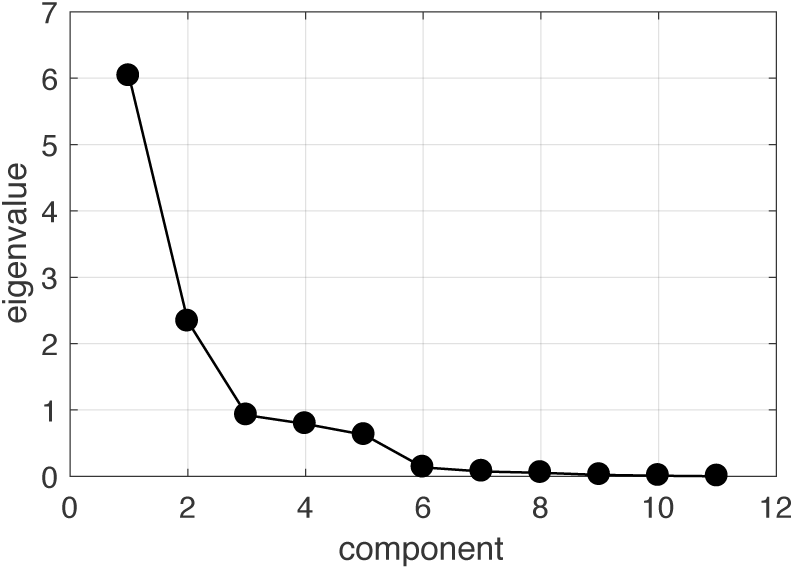
PCA scree plot based on the creativity variables suggests a two-factor solution based on the inflexion point at eigenvalue 3.

To aid in the interpretation of these components, Promax rotation with Kaiser normalization was performed. There was a medium positive correlation between the 2 components (*r* = .33). Components may be correlated as they are both underlying constructs of creativity. Both components show strong loadings on all variables, however, most variables showed substantially loading from only one component (Table 3). This pattern is consistent with a correlated two-component solution, as can also be found in performance vs verbal IQ PCA. Variables related to the number of responses (i.e., count-based scores) showed strong loadings on Component 1 (Self-rating, Fluency, Fluency-valid, Flexibility, Originality, Usefulness); variables related to the quality of the answers showed strong loadings on Component 2 (Examiner-rating, Elaboration, Elaboration-ratio, Originality-ratio, Functionality-ratio). Based on the type of creativity variable each component correlated with most highly, the 2 components may be interpreted as Quantity and Quality as underlying constructs of creativity.

**Table 3:** Pattern matrix for PCA with Promax rotated two-component solution.

|  | Component |  |
| --- | --- | --- |
|  | 1 | 2 |
| Self-rating | **.501** | .220 |
| Examiner-rating | .440 | **.671** |
| Fluency | **1.017** | -.316 |
| Fluency-valid | **1.005** | -.070 |
| Flexibility | **.981** | -.013 |
| Originality | **.794** | .188 |
| Elaboration | .096 | **.832** |
| Usefulness | **.913** | .133 |
| Originality-ratio | .357 | **.506** |
| Elaboration-ratio | -.419 | **.950** |
| Usefulness-ratio | .040 | **.719** |

*Note:* Major loadings are indicated in bold.

**Table 4:**
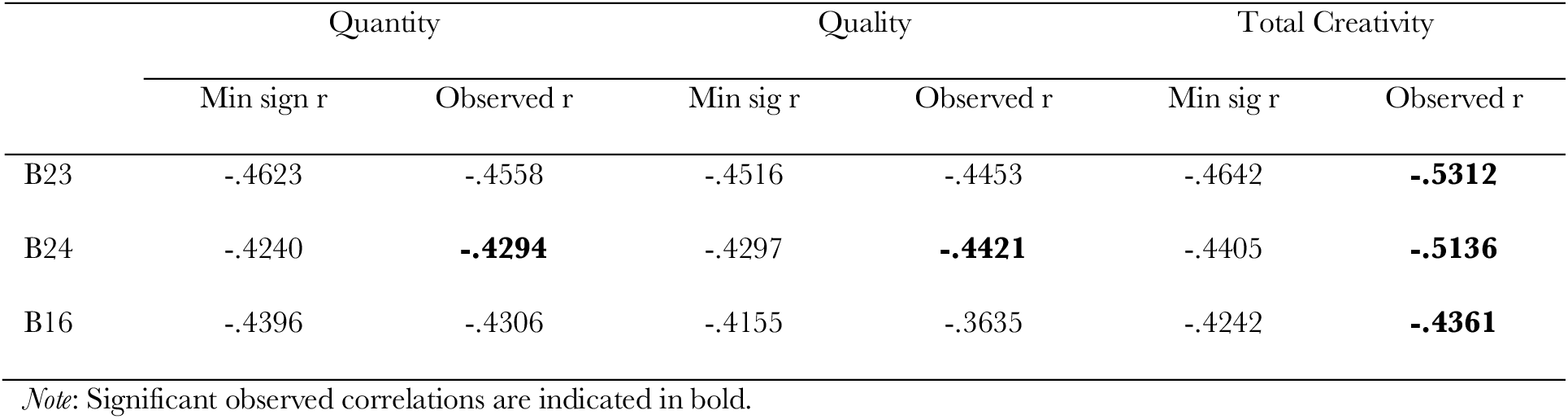
Significance boundaries for minimal correlations (Min sign r) determined by permutation of β-exponents (corrected p < .05, two-sided) and observed correlations for the components Quantity, Quality and Total Creativity for channels B23, B24, B16.

Subsequently, the variables Quantity and Quality with z-transformed component scores were created by means of the regression method. Total Creativity was defined as the sum score of the Quantity and Quality z-scores. Three one-way between-groups analyses of covariance were conducted to investigate whether gender or age had an effect on the variables Quantity, Quality and Total Creativity. The variables were normally distributed and there was no violation of homogeneity of variance between the groups (*p* > .05). There was no significant effect of gender, *F*(1, 37) = 2.49, *p* = .12, partial *η*^2^ = .06, and age, *F*(1, 37) = 0.42, *p* = .52, partial *η*^2^ = .01, for Quantity. This was also applied to the effect of gender, *F*(1, 37) = 2.59, *p* = .12, partial *η*^2^ = .07, and age, *F*(1, 37) = 0.34, *p* = .56, partial *η*^2^ = .01, for Quality. And also for Total Creativity there was no significant effect for gender, *F*(1, 37) = 0.00, *p* = .99, partial *η*^2^ = .00, and age, *F*(1, 37) = 0.53, *p* = .47, partial *η*^2^ = .01.

Age and gender effects were nevertheless used as covariates in all subsequent analyses of brain exponents predicting creativity measures whenever the effect was stronger than p=0.20.

### Power-law characterization of alpha-band amplitude modulation

The average of the exponents *β* across the two resting-state brain activity measurements during the eyes-open resting measurement was used for the analysis. Eyes-open was preferred over the eyes-closed measurement, because dominant alpha activity in the visual brain areas during eyes-closed rest may obfuscates less pronounced alpha oscillatory activity originating in different brain areas. To estimate long-range temporal correlations in ongoing alpha-band oscillations, the standard preprocessing procedure of filtering the EEG from 8.0 –13.0 Hz and extracting the amplitude envelope using the Hilbert transform were used (as in Figure 1 of Smit et al., 2013). Spectral analysis of the amplitude envelope revealed the scaling in the amplitude modulation of alpha oscillations, where *β* is the slope of the straight-line fit in log–log coordinates with average value of .48. The slope represents the decay in temporal correlations. One outlier was observed with average *β*-exponent of 1.27, which was not removed as permutation testing was used for significance.

Figure 3 shows the distribution of average *β*-exponents over the brain, with a minimum value of .32 to a maximum value of .56. The average values for *β* were significantly larger than 0 across all channels in an one-sample t-test, *M* > 0.32, *t*(39) > 7.96, *p* < .001, which suggests that these scaling exponents reflect the presence of long-range temporal correlations. High *β*-exponents have relatively more low-frequency power corresponding to stronger temporal correlations, which reflect more persistent brain states than low exponents (Smit et al., 2013). The effect of gender variated across the 128 channels, abs(*t*(38)) < 2.27, *p* > .03, and age seemed to have no effect on any channel, abs(*t*(38)) < 1.50, *p* > .14. None of these effects were significant after FDR-correction for multiple testing (Benjamini and Hochberg, 1995). A goodness-of-fit of linearity *r*^2^ was estimated in the power-law regression for the spectral analysis. The fit statistic explained more than 75% of the variance across 128 channels of the grand average spectrum.

**Figure 3.**
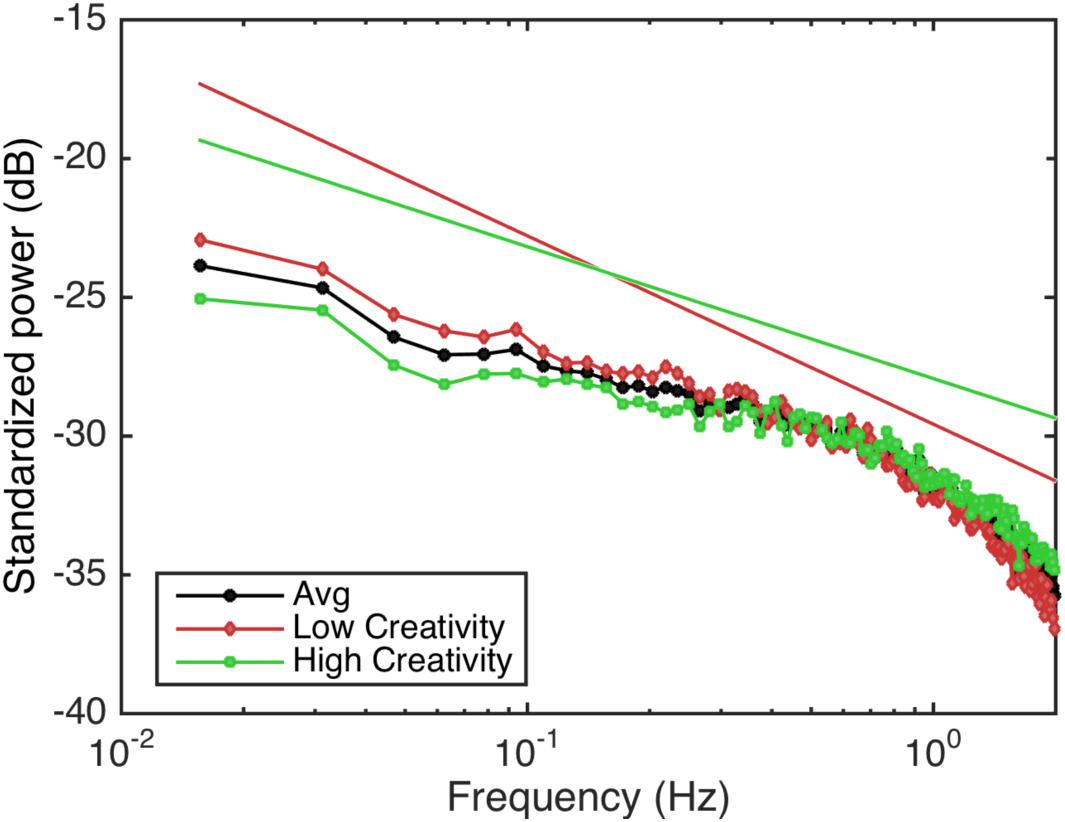
Grand-average power spectra of the amplitude modulation of alpha oscillations at right temporal channel B23 exhibited 1/*f*^*−β*^ scaling as indicated by the log-log linearity for raw spectra <1 Hz. Spectra >2 Hz show a downward curve instigated by the 8-13 Hz filtering (see Smit et al, 2013). Black line is the grand average spectrum (average *β* = 0.48). Red and green are grand average spectra for lower and higher creative ability, respectively, in a median split on Total Creativity. The slope of the line is the scaling exponent and its deviation from zero is a sign of temporal correlations in brain oscillations. The regression lines for lower (*β* = 0.61) and higher (*β* = 0.36) creative individuals are depicted above the spectra for clarity. Excellent linear fit was obtained.

### Low temporal correlations correlates with creativity

We investigated whether the individual variations in power-law frequency scaling exponents of the alpha oscillation modulation could be related to the creativity components Quantity, Quality, and Total Creativity. The distribution of correlations with these creativity components, controlling for gender and age, is shown in Figure 5. The strongest correlations were found over the right temporal areas. These correlations were negative, which means that relatively high exponents (i.e., lower decay in temporal correlation and lower flexibility) are related to low creativity and low exponents (faster decay in temporal correlations and higher flexibility) are related to a high creativity. The variability in scaling exponents between respectively low (*β* = 0.61) and high (*β* = 0.36) creative individuals is shown in Figure 4.

**Figure 4.**
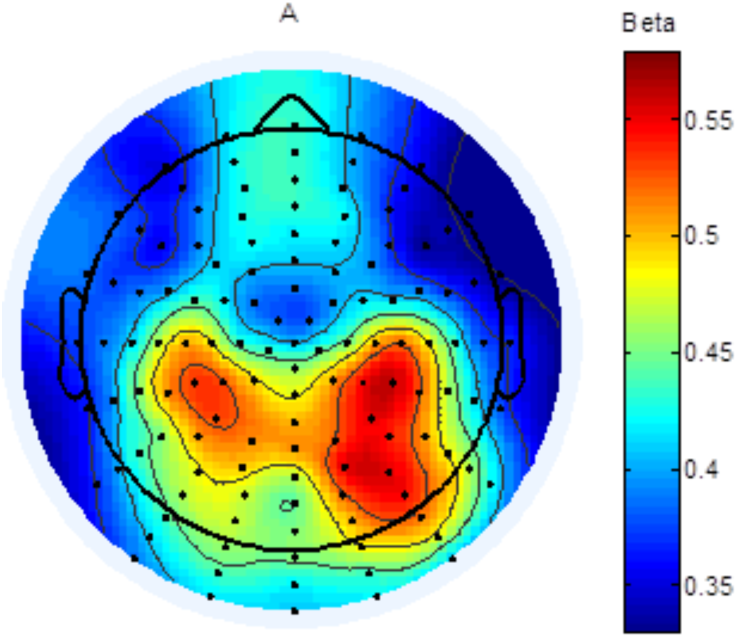
Grand average topographical distribution of the spectral exponents (*β*) are highly consistent with results from Smit et al. 2013, with maximal temporal correlations at right and left parietal signals.

**Figure 5.**
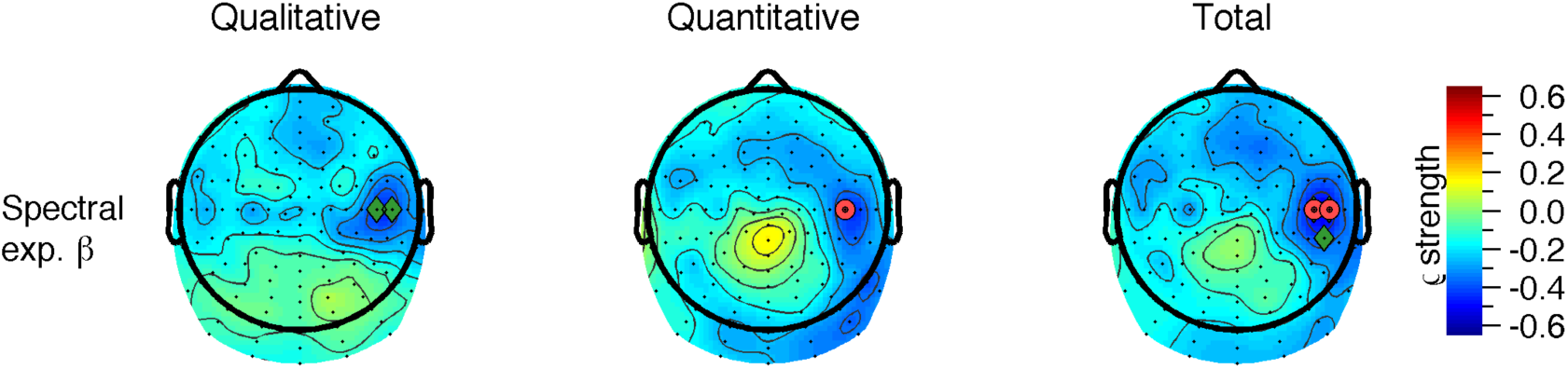
Topographical distribution of the correlations between spectral exponents (*β*), derived from eyes-open resting EEG and Quantity, Quality and Total Creativity. Permutation testing yielded a single significant electrode (p=0.040) for Quantitative Creativity Score (red electrode). For Total Creativity, two electrodes were below the critical minimum correlation (p=0.009 and p=0.012) (red electrodes). Cluster permutation yielded small right temporal cluster that was significant for Qualitative (p=0.035 for two leads) and Total creativity (p=0.034 for three leads; red + green electrodes B23, B24, and B16 of the BioSemi 128 channel layout).

Monte-Carlo Permutation was used to create an empirical null-distribution of the minimum correlation across 128 channels. Note that this test is robust to outliers and correctly handles multiple testing. The test revealed significant correlations coefficients in one channel for Quantity (p=0.040; channel B24 in the BioSemi 128 channel layout) and in two channels for Total Creativity (p=0.009 and p=0.012; channels B24 and B25) over the right temporal areas (Figure 5). We additionally performed a cluster permutation test (Maris and Oostenveld, 2012). This test revealed a two-lead significant cluster for the Qualitative component (channels B24 and B25) and a three lead cluster with Total Creativity (B24, B25, B15), both over right temporal areas (p=0.035 and p=0.034 respectively).

## Discussion

We investigated whether long-range temporal correlations in resting-state alpha oscillations could predict human creativity. Creativity was measured by multiple variables (Self-rating, Examiner-rating and the Alternative Uses Task) and PCA revealed a two (correlated) component structure interpreted as Quantity and Quality. In accordance with the expectation, significant negative correlations were found between the scaling exponents and creativity (for Quantity, Quality and Total Creativity) over the right central brain area. The negative relationship suggests that brains with a tendency for faster decorrelation between brain states—which we hypothesized reflect a more flexible brain—are more creative.

Previous EEG studies of creativity suggested that generation of original ideas is associated with alpha synchronization in the prefrontal cortex (Fink, Swab & Papousek, 2011) and in the parietal and temporal sites of the right hemisphere (Swab et al., 2014). This is partly in line with the results of our research, in which the right central/temporal brain areas appear to be most prominent. The maximal effect at these electrodes may reflect the activity in the sensory motor network which is recruited during execution the AUT (Feng et al., 2019). Using fMRI activity in rest, Feng et al. showed that activity in the bilateral post-central gyri are positively correlated to creative ideation in the alternative uses task. On the network level, they showed evidence for the involvement of several resting-state networks and their integration in creative ideation. In our view, these results complement the current findings, in which the fast state-switching and temporal decorrelation observed over the right central areas could drive the network to quickly switch into a novel organization.

Other neuroimaging studies have implicated the left hemisphere. For example, (Benedek et al., 2014; Dietrich & Kanso, 2010; Fink et al., 2009) revealed strong activation in prefrontal regions and parietal cortex of the left hemisphere during creative ideation. Another fMRI study has found that the right posterior superior temporal sulcus (PSTS) may be selectively involved in verbal creativity, in which verbal creativity reflects the processing of novel metaphoric meanings as well as insight verbal problem solving (Mashal, Faust, Hendler, & Jung-Beeman, 2007). Note that these EEG and fMRI studies used a task-based within-subject design, whereas our current research focused on individual differences by using EEG measurements during eyes-open rest. In addition, none of these studies focused on the temporal dynamics as biomarker of creative ideation. Other literature has suggested a widespread cortical involvement, mainly in prefrontal, temporal, and parietal brain areas (Fink, Swab & Papousek, 2011; Swab et al., 2014).

Although the current results pointed to the involvement of a focused area the right hemisphere, we do not believe that the right central location warrants any firm conclusion about the precise localization. In addition, it may be likely that many other brain areas are involved in the complex behavior elicited by the AUT, but may simply have failed to reach significance due to sample fluctuation. Alternatively, our results may reflect only one aspect in creative ideation, namely the tendency of the brain to switch oscillatory states, which may not be necessary for idea content creation.

Our results seem to contrast with an recent study that reported positive correlations between dwell times in brain states and self-reported openness to experience (Beaty et al., 2018). These authors found that subjects with higher scores on openness—related to creativity—showed longer dwell times in correlational patterns present in resting-state fMRI networks, including the default mode and cognitive control networks. However, these results were obtained using a different methodology, namely, by examining states of consistent cross-connectivity patterns across RSNs, whereas our analysis focused on temporal dynamics or autocorrelation patterns within a single signal. Lower temporal autocorrelations can co-exist with strong cross-regional connectivity. Other research (Simola et al., 2017) reported positive correlations between the DFA exponent of alpha oscillations and cognitive flexibility. However, these authors investigated flexibility in closed-form (convergent) cognitive go-nogo task, which is different from the open-ended AUT measuring divergent creativity.

In conclusion, our results suggest that creativity can be predicted by temporal correlations in resting-state alpha oscillations over right central/temporal cortex. We have found that more creative individuals have lower temporal autocorrelations in functional activity, and thus less persistent brain oscillations than less creative individuals. These individuals show faster state-switching in the alpha oscillations. This flexibility in the intrinsic dynamics of the brain may contribute to an explanation why individuals differ in their behavior and creative abilities, and how innovative ideas and solutions are formed.

## References

Barry, R. J., Clarke, A. R., Johnstone, S. J., Magee, C. A., & Rushby, J. A. (2007). EEG differences between eyes-closed and eyes-open resting conditions. Clinical Neurophysiology, 118, 2765–2773.

Batey, M., & Furnham, A. (2006). Creativity, Intelligence, and Personality: A Critical Review of the Scattered Literature. Genetic, Social, and General Psychology Monographs, 132(4), 355–429.

Beaty, R. E., Benedek, M., Wilkins, R. W., Jauk, E., Fink, A., Silvia, P. J., … Neubauer, A. C. (2014). Creativity and the default network: A functional connectivity analysis of the creative brain at rest. Neuropsychologia. 64, 92–98.

Benedek, M., Jauk, E., Fink, A., Koschutnig, K., Reishofer, G., Ebner, F., Neubauer, A. C. (2014). To create or to recall? Neural mechanisms underlying the generation of creative new ideas. NeuroImage, 88, 125–133.

Cropley, A. J. (2000). Defining and Measuring Creativity: Are Creativity Tests Worth Using?. Roeper Review, 23(2), 72–79.

Dietrich, A., & Kanso, R. (2010). A Review of EEG, ERP, and Neuroimaging Studies of Creativity and Insight. Psychological Bulletin, 136(5), 822–848.

Feng, Q., He, L., Yang, W., Zhang, Y., Wu, X., & Qiu, J. (2019). Verbal Creativity Is Correlated With the Dynamic Reconfiguration of Brain Networks in the Resting State. Frontiers in Psychology, 10.

Fink, A., & Benedek, M. (2014). Review: EEG alpha power and creative ideation. Neuroscience and Biobehavioral Reviews, 44, 111–123.

Fink, A., Grabner, R. H., Benedek, M., Reishofer, G., Hauswirth, V., Fally, M., … Neubauer, A. C. (2009). The Creative Brain: Investigation of Brain Activity During Creative Problem Solving by Means of EEG and fMRI. Human Brain Mapping, 30, 734–748.

Fink, A., & Neubauer, A. C. (2008). Eysenck meets Martindale: The relationship between extraversion and originality from the neuroscientific perspective. Personality and Individual Differences, 44, 299–310.

Fink, A., Swab, D., & Papousek, I. (2011). Sensitivity of EEG upper alpha activity to cognitive and affective creativity interventions. International Journal of Psychophysiology, 82, 233–239.

Furnham, A., & Bachtiar, V. (2008). Personality and intelligence as predictors of creativity. Personality and Individual Differences, 45, 613–617.

Furnham, A., & Nederstrom, M. (2010) Ability, demographic and personality predictors of creativity. Personality and Individual Differences, 48, 957–961.

Gilden, D. L., Thornton, T., Mallon, M. W. (1995). 1/F noise in human cognition. Science, 267,1837–1839.

Guilford, J. P.(1950). Creativity. Am. Psychol., 5, 444–454.

Guilford, J.P. (1967). The Nature of Human Intelligence. New York: McGraw-Hill.

He, B. J. (2014). Scale-free brain activity: past, present, and future. Trends in CognitiveSciences, 18(9), 480–487.

Jauk, E., Benedek, M., & Neubauer, A. C. (2012). Tackling creativity at its roots: Evidence for different patterns of EEG alpha activity related to convergent and divergent modes of task processing. International Journal of Psychophysiology, 84, 219–225.

Jausovec, N., & Jausovec, K. (2000). Differences in Resting EEG Related to Ability. Brain Topography, 12(3), 229–240.

Kefi, S., Rietkerk, M., Roy, M., Franc, A., De Ruiter, P. C., & Pascual, M. (2011). Robustscaling in ecosystems and the meltdown of patch size distributions before extinction. Ecology Letters, 14, 29–35.

Kello, C. T. (2013). Critical Branching Neural Networks. Psychological Review, 120(1), 230–254.

Kello, C. T., Brown, G. D. A., Ferrer-i-Cancho, R., Holden, J. G., Linkenkaer-Hansen, K.,Rhodes, T., &. Van Orden, G. C. (2010). Scaling laws in cognitive sciences. Trends in Cognitive Sciences, 14, 223–232.

Kyaga, S., Lichtenstein, P., Boman, M., Hultman, C., Långstrøm, N., & Landén, M. (2011). Creativity and mental disorder: family study of 300 000 people with severe mental disorder. Br J Psychiatry, 199, 373–379.

Levitin, D. J., Chordia, P., & Menon, V. (2012). Musical rhythm spectra from Bach to Joplin obey a 1/f power law. PNAS, 109(10), 3716–3720.

Linkenkaer-Hansen, K., Monto, K., Rytsala, H., Suominen, K., Isometsa, E., & Kahkonen S. Breakdown of Long-Range Temporal Correlations in Theta Oscillations in Patients with Major Depressive Disorder. The Journal of Neuroscience, 25(44), 10131–10137.

Linkenkaer-Hansen, K., Nikouline, V. V., Palva, J. M., & Ilmoniemi, R. J. (2001). Long-Range Temporal Correlations and Scaling Behavior in Human Brain Oscillations. The Journal of Neuroscience. 21(4), 1370–1377.

Linkenkaer-Hansen, K., Smit, D.J.A., Barkil, A., van Beijsterveldt, T.E.M., Brussaard, A.B., Boomsma, D.I., van Ooyen, A., & De Geus, E.J.C. (2007). Genetic Contributions to Long-Range Temporal Correlations in Ongoing Oscillations. Journal of Neuroscience, 27, 13882–13889.

Mashal, N., Faust, M., Hendler, T., & Jung-Beeman, M. (2007). An fMRI investigation of the neural correlates underlying the processing of novel metaphoric expressions. Brain and Language, 100, 115–126.

Montez, T., Poil, S., Jones, B. F., Manshandenb, I., Verbunt, J. P. A., Van Dijk, B. W., … Linkenkaer-Hansen, K. (2009). Altered temporal correlations in parietal alpha and prefrontal theta oscillations in early-stage Alzheimer disease. PNAS, 106(5), 1614–1619.

Palva, J. M., Zhigalov, A., Hirvonen, J., Korhonen, O., Linkenkaer-Hansen, K., & Palva, S. (2013). Neuronal long-range temporal correlations and avalanche dynamics are correlated with behavioral scaling laws. PNAS, 110(9), 3585–3590.

Post, F. (1994). Creativity and psychopathology. A study of 291 world-famous men. Br J Psychiatry, 165, 22–34.

Power, R. A., Steinberg, S., Bjornsdottir, G., Rietveld, C. A., Abdellaoui, A., Nivard, M. M., … Stefansson, K. (2015). Polygenic risk scores for schizophrenia and bipolar disorder predict creativity. Nature Neuroscience.

Reuter, M., Roth, S., Holve, K., & Hennig, J. (2006). Identification of first candidate genes or creativity: A pilot study. Brain Research, 1069, 190–197.

Runco, M. A., & Acar, S. (2012). Divergent Thinking as an Indicator of Creative Potential. Creativity Research Journal, 24(1), 1–10.

Smit, D. J. A., Linkenkaer-Hansen, K., & de Geus, E. J. C. (2013). Long-Range Temporal Correlations in Resting-State Alpha Oscillations Predict Human Timing-Error Dynamics. The Journal of Neuroscience, 33(27), 11212–11220.

Sternberg, R. J., & Lubart, T. I. (1996). Investing in creativity. Am. Psychologist, 51, 677–688.

Schwab, D., Benedek, M., Papousek, I., Weiss, E. M., & Fink, A. (2014). The time-course of EEG alpha power changes in creative ideation. Frontiers in Human Neuroscience, 8.

Thys, E., Sabbe, B., & De Hert, M. (2012). Creativity and Psychiatric Illness: The Search for a Missing Link – An Historical Context for Current Research. Psychopathology, 46, 136–144.

Torrance, E.P. (1999). Torrance Test of Creative Thinking: Norms and technical manual. Beaconville, IL: Scholastic Testing Services.

Torre, K., Balasubramaniam, R., Rheaume, N., Lemoine, L., & Zelaznik, H. N. (2011) Long-range correlation properties in motor timing are individual and task specific. Psychon Bull Rev, 18, 339–346.

Van Dongen, J., & Boomsma, D. I. (2013). The Evolutionary Paradox and the Missing Heritability of Schizophrenia. Am J Med Genet Part B, 162, 122–136.

Vosburg, S. K. (1998). Mood and the Quantity and Quality of Ideas. Creativity Research Journal, 11(4), 315–324.

